# Maltose-driven carbohydrate status and early raffinose synthesis enhance maturation efficiency in hybrid fir somatic embryogenesis

**DOI:** 10.64898/2026.11.05.764528

**Authors:** Ondřej Skala, Helena Lipavská, Terézia Salaj, Hana Konrádová

## Abstract

Somatic embryogenesis (SE) is a critical biotechnological tool for the large-scale propagation of the *Abies* genus, which faces significant threats from climate change and high commercial demand. While sucrose is a first-choice energy source among conifers, firs often exhibit a dramatic preference for maltose. Using hybrid fir (*Abies alba* x *numidica*), we have shown an order-of-magnitude higher amount of mature cotyledonary embryos cultivated on maltose compared to sucrose. Determination of endogenous carbohydrate balance during maturation and post-maturation phases revealed fundamental difference in spectrum between sucrose (S) and maltose (M) variants, notably massive presence of hexoses in S variant and, remarkably, an earlier accumulation of raffinose in M variant, pointing to establishing a “mature” biochemical profile. After post-maturation desiccation treatment, though both variants accumulated raffinose family oligosaccharides (RFO), their proportion of in total carbohydrate content remained higher in M variant. Based on the results, we propose, that specific features of fir carbohydrate processing machinery (degradation and transport) resulted in slower but steady C and energy supply in M variant and represent the base of maltose superiority.

## 1. Introduction

The *Abies* genus comprises more than 40 ecologically and economically indispensable species. Somatic embryogenesis (SE) offers a powerful asexual propagation method allowing the rapid multiplication of elite genotypes and long-term germplasm storage via cryopreservation (Hazubska-Przybył et al. 2022, Tarraf et al. 2022). Importantly, besides botanical species also embryogenic cultures of interspecific hybrids have been repeatedly derived (e.g., Salajová et al. 1996, Vooková and Kormuťák 2003, 2014). Establishment of an embryonic culture from immature zygotic embryos enables to obtain viable individuals of the crossing in which zygotic embryos abort usually several weeks after pollination (Vooková and Kormuťák 2014).

While SE was first described in *Picea abies* (Chalupa 1985, Hakman and von Arnold 1985), the first successful protocol for the *Abies* genus followed shortly thereafter in 1986 for silver fir (*Abies alba*) (Erdelský and Barančok 1986). Since then, research has successfully expanded to numerous fir genotypes (for review, see Vooková and Kormuťák 2014, Jouini et al. 2025). In line with other conifer models, the process of SE in firs can be divided into four steps: initiation of embryogenic suspensor mass (ESM) from the primary explant (zygotic embryo), ESM proliferation, maturation of somatic embryos and regeneration of plants. However, the technology still faces significant bottlenecks, particularly during the transition from embryo maturation to germination and conversion into plantlet. Even if several of the experimental conditions can be generalised and used for the most *Abies* species, other key ones must be optimised for individual species (Vooková and Kormuťák 2004). Importantly, variables as a carbon source, abscisic acid (ABA) and osmotic agents have been used to increase germination and conversion rates of somatic embryos (e.g. in *A. cephalonica*; Krajňáková et al. 2009). Maltose (usually 3-6%) often gave better maturation response compared to sucrose (Nawrot-Chorabik 2008) and the addition of polyethylene glycol 4000 (PEG, up to 10%) to maturation medium promoted the maturation of somatic embryos (Nørgaard 1997, Salajová et al. 2004, Vooková and Kormuťák 2014). In addition, good germination can be achieved by partial desiccation of cotyledonary somatic embryos (Nørgaard 1997, Vooková et al. 1998, Kvaalen et al. 2005).

The main drawback in evaluating the effectiveness of conifer embryogenic culture modifications is the difficulty in identifying reliable indicators of normal development and the ultimate embryo maturity (Savane et al. 2025; Eliášová et al. 2022). Morphological similarity to zygotic embryos has frequently proven to be a deceptive indicator of physiological maturity and readiness for subsequent development completion (Klimaszewska et al. 2004, Vooková and Kormuťák 2014). Though this approach has been shown to provide some indications of the progress of embryo maturation, it reveals relatively little about the embryo’s developmental potential (Klimaszewska et al. 2004). Consequently, research has shifted toward biochemical markers, such as the accumulation of storage proteins (Kormuťák et al. 2006), lipids, and non-structural carbohydrates like raffinose family oligosaccharides (RFOs; i.e., α-1,6-galactosyl extensions of sucrose) (e.g., Downie and Bewley 2000, Gösslová et al. 2001, Eliášová et al. 2022). Individual carbohydrate levels and their dynamics during embryo maturation was repeatedly suggested as a marker of normal embryo development and metabolic maturity (Lipavská et al. 2000, Iraqi and Tremblay 2001), as well as quantity and quality of reserves in general (Hakman et al. 1990, Attree et al. 1992, Hakman 1993, Misra et al. 1993). Carpenter et al. (2000) suggest that the appropriate marker for the maturity of SE conifers (but above all of the genus *Pinus*) is the level of oligosaccharides of the raffinose family (RFO) and dehydrins, with increased RFO levels resulting in improved sprouting process. But the fundamental difference in the conditions in which zygotic and somatic embryos develop makes assessing the importance of biochemical markers much more complicated. Thus, the precise determination of the role of individual compounds, including non-structural carbohydrates, in the somatic embryogenesis of conifers remains the subject of ongoing research.

In some conifer species, it is necessary to perform desiccation that leads to a reduction in the water content of SEs (sometimes very little) and, importantly, triggers biochemical changes that support further developmental processes: germination and the first steps of plantlet development (Bomal et al. 2002, Eliášová et al. 2022, Savane et al. 2025). The research on the desiccation treatment of conifer SEs began in the 90’s and consisted of partial drying at high relative humidity (Roberts et al. 1990). For *Picea* sp., partial desiccation of SEs is a necessary prerequisite for successful germination (Roberts et al. 1990, Pond et al. 2002), while in larch, on the contrary, desiccation is not required, though it improves synchronisation of germination and conversion to plants (Lelu et al. 1995). In addition, desiccated SEs are also interesting in terms of their long-term storage (Välimäki et al. 2022).

In orthodox conifer seeds, RFO synthesis is natural part of developmental program (Downie and Bewley 2000, Gösslová et al. 2001). Importantly, RFO fulfil a dual role: first, as “starter fuel,” providing rapidly mobilizable energy and carbon skeletons during the earliest hours of imbibition before starch or lipids can be hydrolysed (Downie and Bewley 2000, Savane et al. 2025). Second, and perhaps more critically, they act as osmo- and cryo-protectants (Sengupta et al. 2015). Desiccation of mature somatic embryos repeatedly induced, besides other metabolic changes, the synthesis of RFOs including raffinose, stachyose, and eventually higher members of the family (Konrádová et al. 2003, Businge et al. 2012, Eliášová et al. 2022). RFOs (particularly stachyose) increase cytoplasmic viscosity, disrupt the sucrose crystal lattice, and facilitate a vitrified, “glassy” state, which stabilizes cellular membranes and prevents protein denaturation under water-limited conditions (Caffrey et al. 1988, Bernal-Lugo and Leopold 1995, Hoekstra et al. 2001).

A characteristic feature of SE in the *Abies* genus is its strong preference for maltose, which is rather unique (Nørgaard 1997, Salaj et al. 2004, Jouini et al. 2025). It is hypothesized that the slower hydrolysis of maltose compared to sucrose induces a mild “nutritional stress” that signals the embryo to switch from a proliferative mode to a maturation-focused metabolic program (Scott et al. 1995, Nørgaard 1997). To our knowledge, there are no articles dealing with carbohydrate status dynamics during the fir embryo development (both in the zygotic or the somatic ones). Therefore, this study aims to clarify how the carbohydrate sources modulate carbohydrate metabolism and maturation efficiency in hybrid fir (*Abies alba* x *numidica*).

## 2. Materials and Methods

### 2.1. Plant material

The hybrid fir line AN72 (*Abies alba* x *numidica*), derived from immature zygotic embryo, was maintained in plastic Petri plates (diameter 9 cm) in the dark at 25 °C on modified DCR medium (Gupta and Durzan, 1985) containing 4.4 μM benzylaminopurine, casein hydrolysate (500 mg L^−1^), glutamine (50 mg L^−1^), *myo*-inositol (100 mg L^−1^) and 2% sucrose. The media were solidified with 0.3% Phytagel^TM^ (Sigma-Aldrich). Transfer to fresh media occurred at 3-week intervals.

### 2.2. Maturation and carbohydrate treatments

Maturation was induced by transferring the embryonal suspensor mass (ESM) onto filter paper discs placed on DCR maturation medium containing 10% PEG 4000, casein hydrolysate (500 mg L^−1^), glutamine (50 mg L^−1^), *myo*-inositol (100 mg L^−1^) and 10 μM ABA in 9 cm Petri dishes. Two carbon source variants were compared: maltose variant (M): 3.16% (w/v) maltose·H_2_O; and sucrose variant (S): 3% (w/v) sucrose.

Maturation lasted 8 weeks with one subculture (transfer of the culture together with filter disc) on fresh media with the same composition. This was followed by 1 week of cultivation on ABA and PEG-free media supplemented with the appropriate carbon source (maturation finishing media, FM). Sampling of the material was done always at the end of subcultivation periods: in week 4 (ESM as a whole), in week 8 (isolated embryos, ie) and in maturation finishing phase (ie and mass remaining after somatic embryo isolation, RM).

### 2.3. Post-maturation and germination

Mature SE (9 weeks old) underwent two weeks of high-humidity desiccation at 25 °C in the dark (Eliášová et al. 2022). Following germination took place on DCR medium containing 2% (w/v) sucrose, 1% (w/v) activated carbon (Sigma) and 0.3% (w/v) Phytagel^TM^ in plastic Petri dishes (9 cm diameter). The quantities of the other organic supplements were the same as those in the maturation medium. The first phase of germination took place in the dark at 25 °C. It was terminated once the hypocotyl had elongated and the root had emerged. The cultures were then moved to a cultivation room with a 16-hour photoperiod at 25 °C.

### 2.4. Soluble carbohydrate content determination

Three to six biological replicates of samples (60-200 mg fresh weight) were frozen in liquid nitrogen and stored in −80 °C. Subsequently, the samples were freeze-dried and incubated in 80% methanol at 75 °C for 15 min. Methanol was vacuum-evaporated and the residue re-suspended in ultrapure water before being purified by centrifugation and filtration. The content of soluble carbohydrates was determined using high-performance liquid chromatography (HPLC) (mobile phase water, flow rate 0.5 ml min^-1^, temperature 80 °C) with refractometric detection (Shodex RI-101), Shodex Sugar Column SP0810 (Shodex, Tokio, Japan). Identification of individual carbohydrates was verified using Shodex Sugar Column SC1011 (Shodex, Tokio, Japan). Standard dilutions with known concentrations of sucrose, glucose, fructose, inositol, pinitol, raffinose and verbascose (all Sigma-Aldrich) were used as reference. The HPLC data were evaluated using Clarity 7.8 software (DataApex, Prague, Czech Republic).

### 2.5. Data processing and statistics

Calculations and graphical data processing were performed in Microsoft Office Excel. The data are presented as means ± standard deviations (SD). Statistical data evaluation was performed using NCSS 9 software (NCSS, LLC. Kaysville, Utah, USA). Data were checked for normality and treatment effects (M vs. S variant) were evaluated using standard (equal variance) or Aspin-Welch’s (unequal variance) two-sample *t*-test.

## 3. Results

In this study, we investigated the effect of the type of carbohydrate supplied from the maturation medium (3.16 % maltose or 3% sucrose) on development of the line AN72 of hybrid fir embryogenic culture. The embryo yield and spectrum of endogenous soluble carbohydrates during maturation we examined, first in the entire embryonal suspensor mass (ESM) and subsequently in the isolated developing somatic embryos (SE). The embryos during post-maturation phases were also analysed.

### 3.1. Yield and morphology: the maltose superiority

The carbohydrate added to the maturation medium significantly influenced the culture behaviour. ESM cultured on maltose-containing medium (M variant) was characterized by a slow but steady growth and formation of numerous somatic embryos visible by the end of first subculture. In the case of the culture grown on sucrose-containing medium (S variant), an initial rapid increase in ESM was followed by a marked slowdown in growth and embryo formation was greatly reduced. The ESM often turned brown toward the end of maturation (Fig. 1).

**Figure 1.**
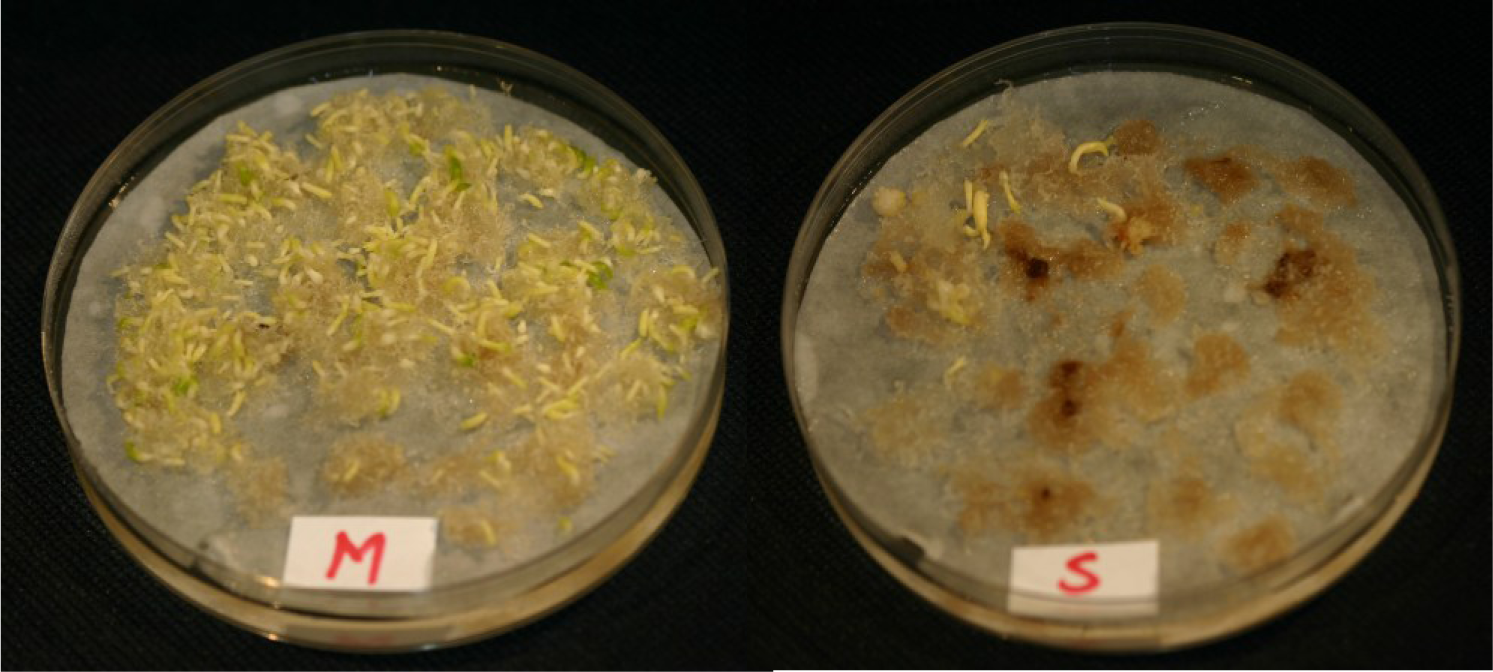
The appearance of the hybrid fir embryogenic cultures grown on the medium with maltose (M variant) and sucrose (S variant) after 8 weeks of maturation and 1 additional week on maturation finishing (MF) medium (for details see section 2.2.).

The carbohydrate source drastically influenced the productivity of the AN72 line. In a single Petri dish (Ø 9 cm), the M variant yielded 133 ± 74 SE, while the S variant produced only 3 ± 2 SE/plate (means ± standard deviations; n=24-27). Morphologically, maltose-grown SE were shorter but robust. Due to the low SE yield in the S variant, some data points regarding the post-maturation stages of development are based on lesser number of samples. Importantly, S variant SE often exhibited impaired development, particularly of the root pole of the embryos, or precocious germination.

### 3.2. Carbohydrate status of S and M variants and maltose induced early raffinose synthesis

Analysis of carbohydrate status (Fig. 2a) revealed a marked decrease in total soluble carbohydrate contents after transfer to the maturation medium in both variants (*t-*test, *p* ≤ 0.01). However, the carbohydrate profiles differed substantially between cultures grown on sucrose and maltose (Fig. 2b): while hexoses significantly predominated in the sucrose variant, the maltose variant spectrum consisted primarily of disaccharides with strongly prevailing maltose. Surprisingly, a considerable content of raffinose was detected in the ESM of M variant as early as in the 4^th^ week of maturation (33.9 μg/mg DW), while, conversely, RFOs remained undetectable in S variant at the same stage. At week 8, the total soluble carbohydrate content was significantly higher in isolated embryos of M variant *(t-*test, *p* = 0.015). Moreover, M variant SE contained 22.2 μg/mg DW of total RFOs compared to 7.2 μg/mg in sucrose-grown SE (significant difference according to *t-*test, *p* ≤ 0.01). As for the other carbohydrates in the spectrum, the situation did not differ significantly: the most abundant carbohydrate in both variants was sucrose, followed by smaller amounts of glucose, fructose, and inositol, with trace amounts of maltose in M variant. After additional week of cultivation on the finishing maturation (MF) medium, the total carbohydrate contents in the isolated embryos did not differ between variants, and no dramatic change was observed when comparing with the situation after 8 weeks of maturation. However, in remaining mass at this stage, the total carbohydrate levels were similar in both variants but significantly higher compared to isolated embryos (*t-*test, *p* ≤ 0.01). Interestingly, the carbohydrate spectra differed markedly between variants: maltose dominated in M variant in contrast to S variant, where spectrum consisted mainly of hexoses with small amount of sucrose. It is worth noting, however, that there was a significant difference in the RFO content, as these carbohydrates were found only in the maltose variant.

**Figure 2.**
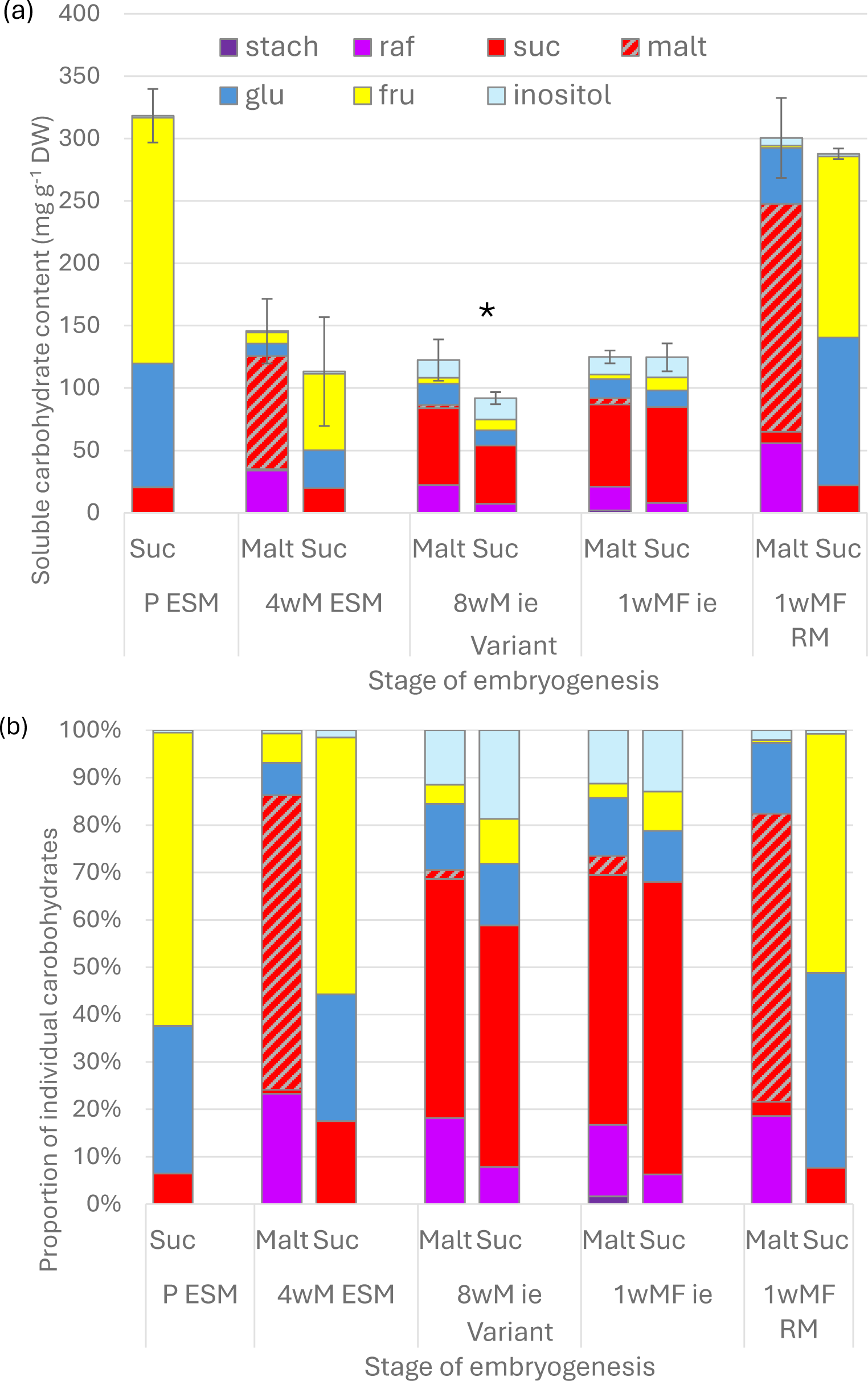
Endogenous soluble carbohydrate contents and spectra in embryogenic line AN72, effect of maltose (Malt) or sucrose (Suc) during maturation. P: proliferation; 4tM, 8tM: 4^th^ and 8^th^ week of maturation; 1tMF: after 1 week of cultivation on maturation-finishing medium. ESM: embryogenic culture analysed as a whole; ie: isolated somatic embryos; RM: mass remaining after embryo isolation. Stach: stachyose; raf: raffinose; glu: glucose; fru: fructose. Total content of soluble carbohydrates (a) and proportion of individual carbohydrates (b). Bars in (a) indicate standard deviations for total carbohydrates, n = 6; for 1tMF Suc n = 3. Asterisk denotes statistical difference in total soluble carbohydrates according to the two-sample *t*-test, *p* ≤ 0.05.

### 3.3. Post-Maturation Carbohydrate Shift

Standard desiccation at high-humidity resulted in a qualitative change in the RFO pool. While raffinose was dominant RFO in isolated SE during late maturation, higher RFO member, stachyose, appeared in S variant and slightly increased *(t-*test, *p* = 0.09) in M variant after the desiccation. The desiccation led to the significant increase in the proportion of RFO in the carbohydrate profile of both variants (*t-*tests, *p* ≤ 0.01; compared with 1t FM of the respective variants). In contrast to the situation observed during maturation, the standard two weeks of slow desiccation led to higher increase of RFO accumulation in SE derived from S variant compared to M variant. Nevertheless, the proportion of RFO in the total carbohydrate content remained higher in the maltose variant (*t*-test, *p* = 0.043). As for the total amount of carbohydrates, the amount found after desiccation was significantly higher in the S variant.

Subsequently, the somatic embryos of both variants were transferred to germination media. Figure 3 shows 3-week-old fir seedlings; the SE of M variant did not exhibit normal development instead we observed abnormalities in them, particularly poorly developed roots. In both variants, the endogenous carbohydrate status of harvested seedlings was characterized by the complete disappearance of RFO and presence of considerable proportion of sequoitol along with lower amount of pinitol (both are *O*-methyl-inositol isomers). Nevertheless, M variant contains significantly higher sum of methyl-inositols compared to S variant (*t*-test, *p* = 0.030).

**Figure 3.**
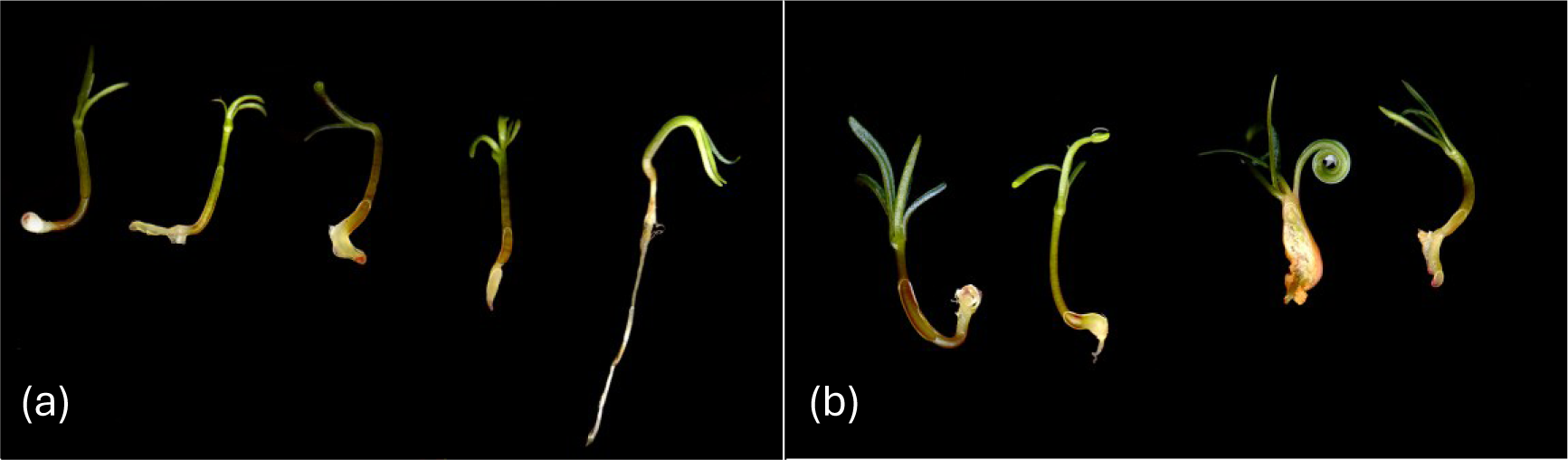
Appearance of 3-week-old germinating seedlings (emblings) matured on medium with maltose (a) or sucrose (b).

**Figure 4.**
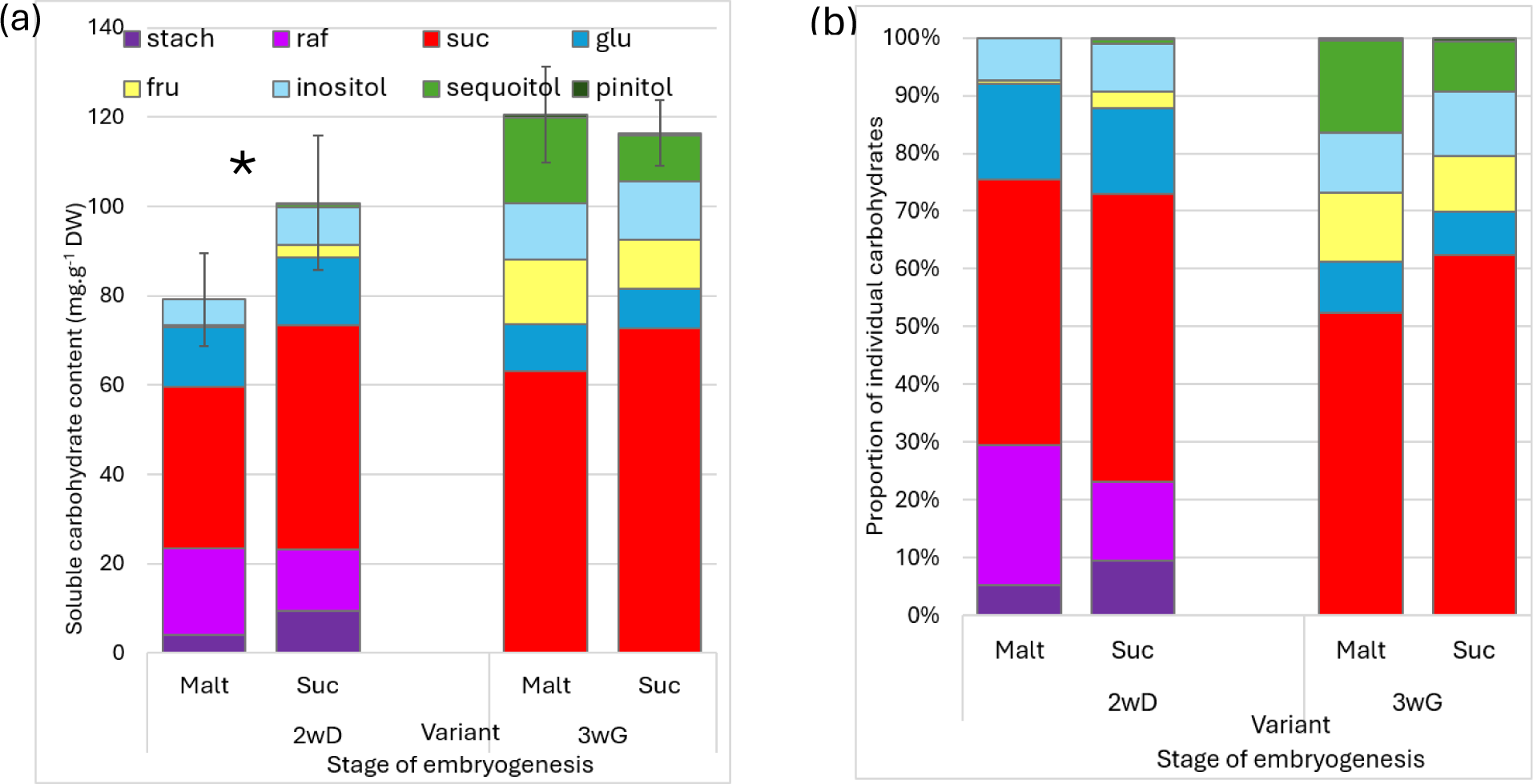
Endogenous soluble carbohydrate contents and spectra in embryogenic line AN72, effect of maltose (Malt) or sucrose (Suc) after 2 weeks of desiccation at high humidity and subsequent 3 weeks of germination. Isolated embryos; 2wD: 2 weeks of desiccation; 3wG: 3 weeks of germination. Stach: stachyose; raf: raffinose; glu: glucose; fru: fructose. Total content of soluble carbohydrates (a) and proportion of individual carbohydrates (b), n = 5. Bars in (a) indicate standard deviations for total carbohydrates. Asterisk denotes statistical difference in total carbohydrates according to the two-sample *t*-test, *p* ≤ 0.05.

## 4. Discussion

In fir somatic embryogenesis, numerous studies have demonstrated frequent maltose superiority to support maturation efficiency. However, to our knowledge, there is no sound experimental basis for explaining this phenomenon. In spruce, the dynamics of the endogenous carbohydrate content proved to be a good indicator of favourable embryo development when evaluating the effects of protocol modifications (Lipavská and Konrádová 2004, Businge et al. 2014, Hudec et al. 2016, Eliášová et al. 2022). In this study, we made use of this experience and tried to reveal the physiological basis of the phenomenon by monitoring endogenous carbohydrate dynamics as affected by the exchange of exogenous carbohydrate supply. Surprisingly, there is a complete lack of information on carbohydrate status during the somatic embryogenesis of the fir.

### 4.1. Maltose as a yield-enhancing factor acting through nutritional modification

With the use of hybrid fir embryogenic line AN72 (*Abies alba* x *numidica)*; Salaj et al. 2004) we confirmed the dramatic disparity in embryo yield between the application of maltose. The advantage of using this line stems from the possibility to perform analysis even on sucrose-supported embryos, though the yield of mature embryos was very low. The data indicated that sucrosés failure to promote high-quality maturation in line AN72 likely resulted from its rapid hydrolysis by wall invertases in the apoplast, which led to a surplus of hexoses (glucose and fructose) and potentially induced hypoxia or ethanol toxicity (Nørgaard 1997, Iraqi et al. 2005). In the early stages of maturation (after 4 weeks on maturation medium), the culture was rich in hexoses, particularly fructose, which was probably due to the high activity of hexose transporters and also the lower activity of enzymes capable of o processing fructose in the cytosol. The pronounced accumulation of fructose in our sucrose-grown cultures, accompanied by tissue browning and impaired embryo development, closely mirrors the fructose-induced developmental block reported in *Picea abies* (Businge et al. 2012, Businge and Egertsdotter 2014). Fructose is a highly reactive reducing keto-sugar making it an extraordinarily potent initiator of non-enzymatic glycation via the Maillard reaction (Suarez et al. 1989, Semchyshyn et al. 2011). Fructose accumulation therefore triggers protein carbonylation, DNA damage, and glutathione depletion, ultimately causing tissue necrosis and cellular death (Baynes 2002, Businge and Egertsdotter 2014, Tola et al. 2021). Thus, the low quality and quantity of embryos in this variant can be attributed to fructose toxicity and associated oxidative stress. This is further consistent with the finding that browning in *Picea abies* was caused not by the accumulation of phenolic compounds, but rather by their increased oxidation in an environment with a reduced content of low-molecular-weight ROS scavengers (Švorc Štepánová et al. 2026). Moreover, high level of fructose was reported to down-regulate the expression of *WUS/WOX*, a stem cell-promoting transcription factor, via the fructose activated sensor FINS1 (fructose 1,6-bisphosphatase) which acts as a bifunctional factor (Li et al. 2023). On top of all that, hexoses can interact with abscisic acid (ABA), auxin, and ethylene pathways to modulate cell division and development (Li et al. 2023).

In contrast, the slower utilization of exogenously provided maltose very likely ensures a moderate, but more stable, carbon supply to nourish maturing embryos. We can base this assumption on the observed high proportion of maltose in the carbohydrate spectrum of ESM after 4 weeks of maturation, as well as in the residual mass remaining after the isolation of embryos upon finishing maturation. We suggest that this is the result of very inefficient apoplastic degradation of maltose, and low ability of transporters to help maltose to cross cell membranes. Together it might mimic the slow-release nutritional environment of a developing seed (Scott et al. 1995, Nørgaard 1997). This probably brought about only mild and balanced „nutritional stress” which acts as a biochemical signal, steering the tissue away from simple proliferation toward the SE maturation program, including the early synthesis of RFO, with capacity to act as protective compounds (Scott et al. 1995, Nørgaard 1997, Sengupta et al. 2015).

### 4.2. RFO accumulation: a developmental program vs. stress response

Another striking difference between the two treatments was the presence of raffinose in the ESM after 4 weeks of maturation exclusively in the maltose variant. If we consider that RFO synthesis is often associated with stress exposure and the stress load is due to high endogenous hexoses levels higher in sucrose variant, then the stress-based idea faces a problem. However, it is necessary to keep in mind that the synthesis of RFO is a standard part of developmental program of conifers, and thus we have to search for another signal that strengthens or triggers embryonic development ((Downie and Bewley 2000, Gösslová et al. 2001). This assumption is reinforced by the finding that the few embryos that formed on sucrose medium also synthesized RFO in later stages of development (after 8 weeks of maturation) even though the amount of RFO remained higher in maltose supported ones. It should be noted that the carbohydrate profiles were already similar in both variants at this stage of development, with sucrose clearly predominating; sucrose apparently originated from resynthesis even in the sucrose variant (Kubeš et al. 2014). We can further speculate that the individual components of the carbohydrate spectrum, specific in sucrose and maltose variants, especially at the early stages of maturation, have entirely different signalling potentials for the plant cells. Therefore, we propose that the carbohydrate signalling background established in maltose variant may be the key factor not only for RFO synthesis but also for promoting embryo formation.

In this context, it may be interesting to note that the carbohydrate status fundamentally differs from the Norway spruce model (*Picea abies*), where RFOs are almost entirely absent until triggered by post-maturation stressors like cold or drying (Gösslová et al. 2001, Konrádová et al. 2003, Businge et al. 2013, Eliášová et al. 2022). On the other hand, there is an interesting similarity between these two conifer models: as maturation progresses, the proportion of endogenous sucrose in the carbohydrate profile increases in both species, even in fir cultures grown on maltose. In fir, the early detection of raffinose in the ESM by week 4 suggests that RFO synthesis is a part of the developmental program and indicates the embryos’ readiness for further development, which is triggered, in this case, by maltose signalling. In *Picea*, even synergistic stressors have to be applied to reach peak RFO levels, whereas in *Abies*, the metabolic machinery for RFO production is already activated while the embryos are still in contact with the high-water-potential maturation medium. This highlights that RFO metabolism in conifers is strictly species-specific, and no general model for “biochemical maturity” can be applied across the Pinaceae (Lipavská and Konrádová 2004, Jouini et al. 2025, Stojičić et al. 2026).

### 4.3. Post maturation embryo behaviour as affected by maltose supported maturation

The observed synthesis of RFO during maturation raised the question of what effect the application of a post-maturation desiccation treatment might have on the subsequent content and spectrum of carbohydrates. The desiccation exposure (more precisely high-humidity treatment) significantly enhanced the proportion of RFO in both variants. Moreover, besides raffinose, the stachyose proportion also increased in the maltose variant and stachyose appeared in sucrose one as well. To our knowledge, in fir, there is no information on carbohydrate balance even during post-maturation phases. However, desiccation treatment is frequently reported as beneficial for further successful development (Nørgaard 1997, Vooková and Kormuťák 2003, 2014, Salaj et al. 2020, Jouini et al. 2025). In spruce, where desiccation treatment is critical condition for embryo germination, it is accompanied by RFO synthesis, including stachyose (Gösslová et al. 2001, Konrádová et al. 2003, Eliášová et al. 2022). Stachyose is reported to be more effective than raffinose at increasing cytoplasmic viscosity and raising the transition temperature for the vitrified glassy state (Bernal-Lugo and Leopold 1995, Sun 1997, Buitink and Leprince 2008, Sengupta et al. 2015). Based on data presented here, it is unfortunately not possible to establish a clear relationship between the beneficial effect of desiccation and RFO accumulation in fir embryogenesis. This is further illuminated by recent transcriptomic findings in desiccation-treated conifer embryos showing that RFO synthesis (via PaGOLS and PaRFS) is co-regulated with cell wall remodelling under PaMYB12 transcription factor control (Ling et al. 2025, Savane et al. 2025). Anyway, our previous data alongside broader literature - even in the case of well-studied spruce model – clearly show that this relationship is more complex, as the induction of RFO synthesis e.g. by cold, though beneficial, proved to be insufficient on their own to guarantee high-quality germination (Konrádová et al. 2003, Businge et al. 2013, Eliášová et al. 2022, Välimäki et al. 2022). During germination, the RFO fraction disappeared completely from the spectrum, suggesting that this carbohydrate plays a role in promoting germination as an immediately mobilizable carbon and energy source (Downie and Bewley 2000, Gangl and Tenhaken 2016). Interestingly, differences in the carbohydrate spectrum caused by maltose versus sucrose supply in maturation can even be traced up to the germination phase, particularly a significantly higher proportion of *O-*methylated inositol derivatives in the embryos of the maltose variant. However, the significance of this difference, if any, remains unclear.

Recent findings suggest that the environmental conditions during SE - such as temperature or osmotic stress - can induce an epigenetic memory that influences the future phenology and physiology of the emblings (Lobo et al. 2022; Trontin et al. 2025). We propose that similarly, maltose as a carbon source in fir maturation likely influences the “metabolic memory” of the embryo. While our study has focused on the carbohydrate markers of this process, the broader implication is that optimizing the maturation environment not only improves yield but also defines the future fitness and adaptive potential of the resulting somatic plants in forest environments.

## Conclusions

- Maltose is the essential driver for hybrid fir (*Abies alba* x *numidica*) SE, ensuring massive productivity of heathy embryos.
- The endogenous carbohydrate profiles of the maltose- and sucrose-supported variants differed significantly: low hexose levels and the early synthesis of raffinose were most noteworthy characteristic of the maltose variant, indicating a distinct signalling network status.
- Desiccation enhanced total RFO accumulation in both variants, while stimulating stachyose synthesis; germination caused complete RFO consumption. Notably, the carbohydrate profile differences between variants, manifested in their *O*-methyl-inositol derivative levels during germination) persisted until the end of the monitored phase.
- The generic application of conifer sucrose-based maturation strategies is incompatible with the unique metabolic requirements of particular *Abies* species.

## Acknowledgements

This work is based upon work from COST Action CA21157 COPYTREE (European Network for Innovative Woody Plant Cloning), COST (European Cooperation in Science and Technology).

## Authors’ Contributions

Ondřej Skala: Investigation, Visualization, Formal analysis. Helena Lipavská: Writing – review and editing, Methodology, Supervision. Terézia Salaj: Writing – review and editing, Conceptualization, Methodology. Hana Konrádová: Writing - original draft, Writing – review and editing, Data curation, Project administration, Conceptualization. All authors read and approved the final manuscript.

## Data and Materials Availability

The data generated during the study are available from the corresponding author on reasonable request.

